# Predicting Differentially Methylated Cytosines in TET and DNMT3 Knockout Mutants via a Large Language Model

**DOI:** 10.1101/2024.05.02.592257

**Authors:** Saleh Sereshki, Stefano Lonardi

## Abstract

DNA cytosine methylation is an epigenetic marker which regulates many cellular processes. Mammalian genomes typically maintain consistent methylation patterns over time, except in specific regulatory regions like promoters and certain types of enhancers. The dynamics of DNA methylation is controlled by a complex cellular machinery, in which the enzymes DNMT3 and TET play a major role. This study explores the identification of differentially methylated cytosines (DMCs) in TET and DNMT3 knockout mutants in mice and human embryonic stem cells. We investigate (i) whether a large language model can be trained to recognize DMCs in human and mouse from the sequence surrounding the cytosine of interest, (ii) whether a classifier trained on human knockout data can predict DMCs in the mouse genome (and vice versa), (iii) whether a classifier trained on DNMT3 knockout can predict DMCs for TET knockout (and vice versa). Our study identifies statistically significant motifs associated with the prediction of DMCs each mutant, casting a new light on the understanding of DNA methylation dynamics in stem cells. Our software tool is available at https://github.com/ucrbioinfo/dmc_prediction.

## Introduction

DNA methylation is an epigenetic marker which directly or indirectly regulates several critical cellular processes, including gene expression, genome stability, transposon suppression, and gene imprinting (see, e.g., (14; 15; 16; 17; 18; 19; 20; 21; 22; 23; 24; 25; 26)). The most common form of DNA methylation, known as 5-methylcytosine (5mC), involves the attachment of a methyl group to the fifth carbon of a cytosine residue. Abnormal methylation patterns in humans have been associated with diseases, including cancer and imprinting syndromes (see, e.g., (29; 30; 31; 32)).

In mammals, DNA methylation primarily occurs in CpG dinucleotides, with most of them being methylated (28). Mammalian genomes typically maintain consistent CpG methylation patterns over time, except in specific regulatory regions like promoters and certain types of enhancers (27). In these variable regions, the dynamics of methylation and demethylation is orchestrated by a complex cellular machinery, in which the enzymes DNMT3 (A/B) and TET play a major role. DNMT3A and DNMT3B are DNA methyltransferases that can add a new methyl group to cytosines, e.g., during development and cellular differentiation (34; 33). TET is an enzyme that catalyzes the conversion of 5-methylcytosine to 5-hydroxymethylcytosine and its oxidized derivatives. The conversion of 5-hydroxymethylcytosine and its derivatives ultimately leads to active DNA demethylation (35).

Knockout experiments that disrupt DNMT3 and TET have allowed life scientists to unravel the complex dynamics of DNA methylation changes over time and space, and across cell types. During pluripotency stages, DNMT3 and TET modulate the epigenetic landscape, thus influencing cellular differentiation (36; 2). During post-fertilization reprogramming, the embryo undergoes a two-phase process in which it loses gamete-specific DNA methylation patterns inherited from the oocyte and sperm, with the initial active demethylation of the paternal genome by TET3 followed by subsequent passive dilution of DNA methylation during cell divisions (37).

TET and DNMT3s are crucial in regulating fetal organ development and tissue generation, through DNA methylation and histone modifications (55; 54; 56; 57; 58). Their dysregulation is linked to human diseases, particularly cancers (59; 60; 61; 62; 63). Although the importance of TETs is well recognized, their precise mechanisms of action is not well understood. Several studies have shown that DNMT3 and TET, both individually and in combination, influence DNA methylation patterns in human embryonic stem cell lines (e.g., (2)). Chao *et al*. also studied the interactions between TET1, DNMT3A, and DNMT3B in human embryonic stem cells, and how these interactions collectively influence global methylation patterns (9).

Given the importance of DNMT3 and TET in developmental biology and embryogenesis, there is strong interest in characterizing which cytosines are affected by these two classes of enzymes. A few recent studies attempted to capture the sequence preference for DNMT3 and TET. For instance, in (39) the authors showed that TET has a sequence preference for CG dinucleotides within specific transcription factor-binding sites, indicating that its activity in catalyzing DNA demethylation is influenced by the underlying sequence context. The study in (9) also reported that TET1 prefers binding to specific genomic regions. This appears to be also true for DNMT3. Jeltsch et al. (40) demonstrated that the enzymatic activities of DNMT3A and DNMT3B are influenced by the sequence context of their target sites. As part of these studies, several DNMT3 and TET knockout methylation data sets for human and mouse have been produced (see, e.g., (2; 4; 6)). These data sets open the possibility to investigate whether one could predict which specific cytosines are affected by DNMT3 and TET using a machine learning model.

Here we explore for the first time the problem of predicting differentially methylated cytosines (DMC) in TET and DNMT3 knockout mutants, using exclusively the underlying DNA sequence around the cytosines. Our predictor, called L-MAP (Language model-based Methyltransferases Activity Predictor), is transformer-based large language model that utilizes contextual sequence information to predict the enzymatic activity of DNMT3 and TET on cytosines.

We envision the main use of L-MAP as a tool to impute missing and/or uncertain DMCs obtained from wet lab experiments. In this study, we also investigate (1) whether training L-MAP on DNMT3 knockouts can be used to predict TET activities, and vice versa; (2) whether training L-MAP on human knockout data can be used to predict enzymatic activity on mice, and vice versa; (3) whether the methylation levels of nearby cytosines can help L-MAP predicting DMCs with higher accuracy; (4) whether L-MAP has learned sequence motifs known to be associated with the activity of DNMT3 and TET enzymes.

A deeper understanding of cell functions can lead to significant advancements in medical research, therapeutic development, disease prevention, and diagnostic techniques, such as drug discoveries (64; 65; 66). Some studies have identified interacting partners for TETs and DNMT3s (67; 68; 69; 70). Here, we have identified transcription factor binding site (TFBS) motifs that may be linked to TET and DNMT3 activity in pluripotent cells. These findings can open new avenues for understanding the functions of these methyltransferases and lead to advancement in treatment strategies and novel drug discoveries.

## Results

Seven data sets, across three studies, were used to train and test L-MAP. In the first study by Charlton *et al*. (2), the authors utilized CRISPR-Cas9 to create an array of gene knockouts in human embryonic stem cells (ESC) involving DNMT3s and TETs, both individually and in combination. The following knockout configurations were established: (i) in the DNMT3KO ESC line, both DNMT3A and DNMT3B genes were deactivated; (ii) in the TETKO ESC line, TET1, TET2, and TET3 genes were knocked out; (ii) in the QKO ESC line, TET1, TET2, TET3 and DNMT3B were deactivated; and (iv) in the PKO ESC line, TET1, TET2, TET3, DNMT3A and DNMT3B were knocked out. The second study by Gu *et al*. (4) involved DNMT3A and DNMT3B knockout in mouse ESC. The third study by Ansari *et al*. (6) involved TET2 and TET3 knockout in mouse intestinal stem cells (ISC).

All ten datasets (seven knockout and three wild type) (i) were obtained using whole genome bisulfite sequencing using Illumina sequencing instruments and (ii) were processed using the BSMAP pipeline (1) for mapping bisulfite-treated reads to the reference genome. In our experiments, we used the methylation levels provided by the authors. However, to ensure that we could compare methylation level across different studies, we re-analyzed the three wild-type samples using a common software pipeline. We processed the three sets of Illumina reads through Bismark (5) using default parameters. The methylation levels obtained from our pipeline matched almost exactly the methylation levels provided by the authors: the mean square difference between our levels and those provided by the authors were less than 2%.

Supplemental Figure 2 reports the genome-wide methylation levels for the three wild type and seven knockout data sets. In general, the methylation level of a cytosine *c* ranges from 0 to 1, where 0 indicates that none the cells in the sample are methylated on *c*, and 1 indicates that all the cells in the sample are methylated on *c*. Observe that the average methylation level is in the range 0.7–0.8 for all data sets, except for the DNMT3A knockout data set on the mouse ESCs.

The methylation levels for the seven knockout and three wild type datasets were used to determined seven sets of differentially methylated cytosines (DMC). A cytosine was determined to be differentially methylated when its methylation level for the knockout was significantly higher or lower than its methylation level in the wild type (details in Methods). Supplemental Figure 1 reports the number of differentially methylated cytosines on the seven data sets. Observe that the number of differentially methylated cytosines ranges from about 100 thousand in the DNMT3B knockout dataset for mouse ESC, to about 1.5 million in the DNMT3A knockout for mouse ESC. Based on this, we chose a sample size of 100 thousand cytosines for each data sets, half of which were differentially methylated (and the other half was not). The sample included 100 thousand 512 bp-long DNA sequence centered around the chosen cytosines, along with the corresponding binary label (1 indicated a DMC, 0 otherwise). We evaluated the impact of size of the training dataset on L-MAP’s accuracy in Supplemental Figure 7. Observe that the accuracy improves up to a sample size of 100,000. Further increases in the sample size do not significantly improve L-MAP’s accuracy.

L-MAP was trained on 90% of the cytosines (chosen uniformly at random from each data set) and tested it on the remaining 10%. To ensure consistent results across different random train-test splits, we computed the variance of L-MAP’s accuracy across five random samples of the training set for TETKO and DNMT3KO. The average and standard deviation for L-MAP’s accuracy is illustrated in Supplemental Figure 5. Observe that the deviation on L-MAP’s accuracy is very small across different random samples for the training set, which allowed us to rely on the results of a single run for the rest of the experiments.

Figure 1-D shows the methylation levels of human ESC wild-type and TET knockout cytosines in the region [2493500,2497500] of chromosome 19, as a qualitatively example of the training data. The middle panel shows the difference in methylation level between the two cell lines, where the red dots indicate differentially methylation cytosines (blue otherwise). The lower panel shows L-MAP’s predictions of differentially methylation cytosines based on the sequence context around the cytosine. Observe how L-MAP makes accurate predictions in the middle portion of this region.

**Fig. 1.**
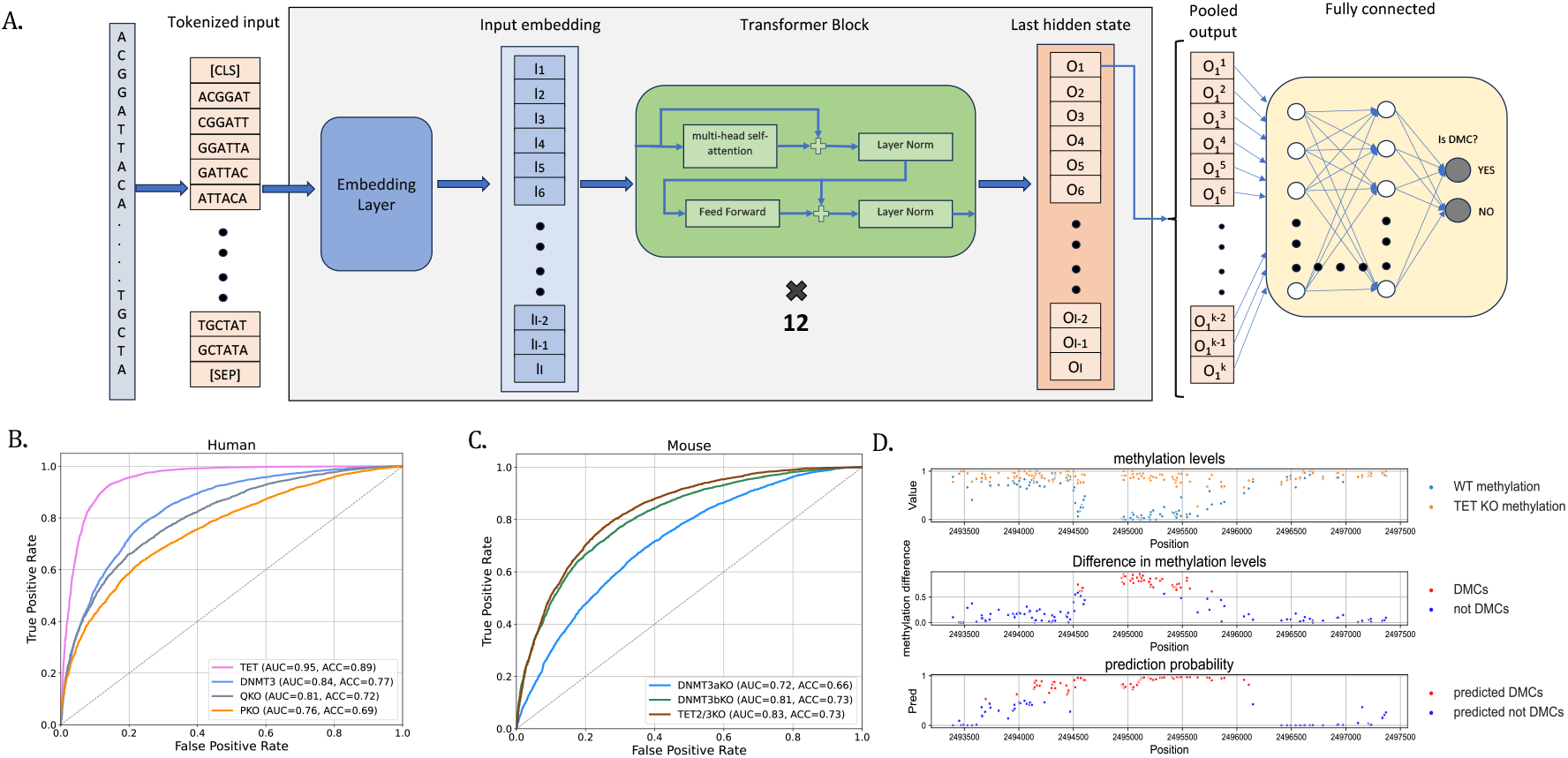
**(A)** the architecture L-MAP; **(B)** ROC curves for the performance of L-MAP on human TET and DNMT3 knockout datasets (AUC=area under the curve, ACC=accuracy); **(C)** ROC curves for the performance of L-MAP on mouse TET and DNMT3 knockout datasets (AUC=area under the curve, ACC=accuracy); **(D -upper panel)** methylation levels of human ESC wild-type and TET knockout cells in the region [2493500,2497500] of chromosome 19; **(D -middle panel)** the difference in methylation level between wild-type and knockout, red dots indicate differentially methylation cytosines; **(D -lower panel)** predictions generated by L-MAP based on contextual sequence information, red dots indicate cytosine that are predicted to be differentially methylated

Figure 1-B show the ROC curves for the binary classification performance of L-MAP on the four human knockout data set. Observe that the best classification performance was achieved on the TETKO dataset in which TET1, TET2, and TET3 genes were knocked out (area under the curve 0.95, accuracy 0.89). The second best was on the DNMT3KO dataset, in which both DNMT3A and DNMT3B genes were knocked out. The quadruple knockout (QKO) and quintuple knockout (PKO) had lower accuracy and AUC compared to TETKO and DNMT3KO. Our hypothesis is that mixing multiple enzymatic knockouts in QKO and PKO makes it harder for the classifier to capture their sequence specificity. However, the fact that L-MAP can still classify differentially methylated cytosines in the QKO and PKO suggests the existence of common sequence signatures between the two classes of enzymes.

Figure 1-C shows the ROC curves for the binary classification performance of L-MAP on the three mouse knockout data set. Again, observe that the best classification performance was achieved on the TET2/3KO dataset in which both TET2 and TET3 genes were knocked out (area under the curve 0.83, accuracy 0.73). These results in human and mouse suggest that the TET activity is more sequence-dependent than DNMT3. The performance of L-MAP on the DNMT3bKO dataset was the second best.

### Cross knockout prediction

In the following experiments we carried out a set of cross-knockout and cross-species predictions. In one set of experiments, L-MAP was trained on one knockout dataset and tested on a different knockout enzyme. In the second set, L-MAP was trained on human knockout data, and tested on mouse data, or vice versa.

The cross-species L-MAP’s accuracy is visualized in Figure 2-A and Figure 2-B, for human and mouse, respectively. Observe that in most cases the highest accuracy is observed when L-MAP is trained and tested on the same data set, as expected. However, there are some exceptions. L-MAP’s accuracy is higher when trained on human PKO and QKO data sets and tested on TET data sets, compared to being tested on the same knockout dataset. This can be explained by the presence of shared patterns in PKO and QKO cell lines, both of which include the knockout of TET.

**Fig. 2.**
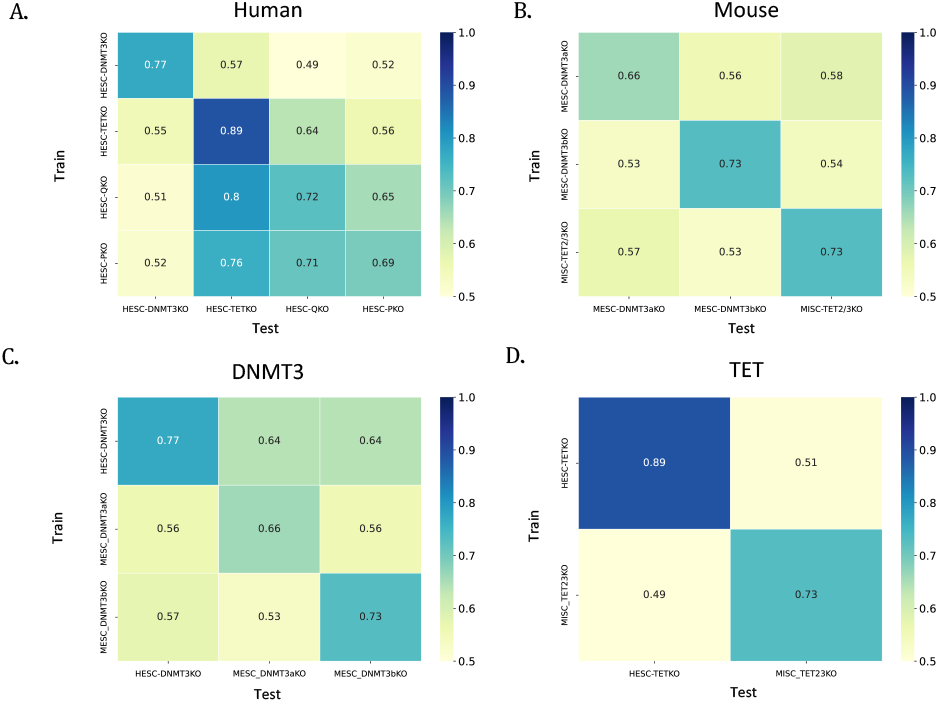
**(A)** L-MAP’s accuracy when trained on a human knockout dataset and tested on another human knockout dataset **(B)** L-MAP’s accuracy when trained on one mouse knockout dataset and tested on another mouse knockout dataset **(C-D)** L-MAP’s accuracy when trained on a human (mouse) knockout dataset and tested on a mouse (human) knockout datasets

The cross-knockout L-MAP’s accuracy is illustrated in Figure 2-C and Figure 2-D. Figure 2-C reports the results on three data sets: two for mouse ESC (DNMT3AKO and DNMT3BKO) and one for human ESC (DNMT3KO, which includes DNMT3A and DNMT3B knockout). Figure 2-D reports the results on two data sets: one for human ESC (TETKO, corresponding to TET1, TET2, and TET3 knockout) and one for mouse ISC (TET23KO, representing TET2 and TET3 knockout). Observe again that the highest accuracy is achieved when the model is trained and tested on the same dataset. Also observe that in the case of TET the cross-species experiment yields significantly lower accuracy, suggesting that the underlying sequence contexts associated with TET activity are likely to be different in the two species.

### Motif analysis

The objective of this analysis was to extract “knowledge” from the LLM to gain insights on the sequence context employed by L-MAP to make predictions about DMCs. Briefly, we used the attention layer of L-MAP to identify DNA sequences associated with DMCs and DNA sequences associated with non-DMCs. These positive and negative examples were processed using STREME (41), to obtain motifs and corresponding p-values (see Methods for details). Figure 3 reports the motifs with the lowest p-value for each of the seven knockout datasets (the lowest three p-value motifs are reported in Supplemental Figures 8 and 9). We utilized JASPAR (38) to search for known motifs that matched our motifs. The best matches are reported in last four columns of Figure 3. First observe that all the motifs belong to the C2H2 zinc finger factors, which are known to have a role in methylation and demethylation processes (see, e.g., (11; 12; 13; 3)). For instance, zinc finger protein ZNF615 plays a significant role in embryonic stem cell development through DNA methylation by facilitating the recruitment of DNA methyltransferases to specific genomic regions (42). Most of the matching motif are also associated with molecular mechanisms in embryonic stem cells.

**Fig. 3.**
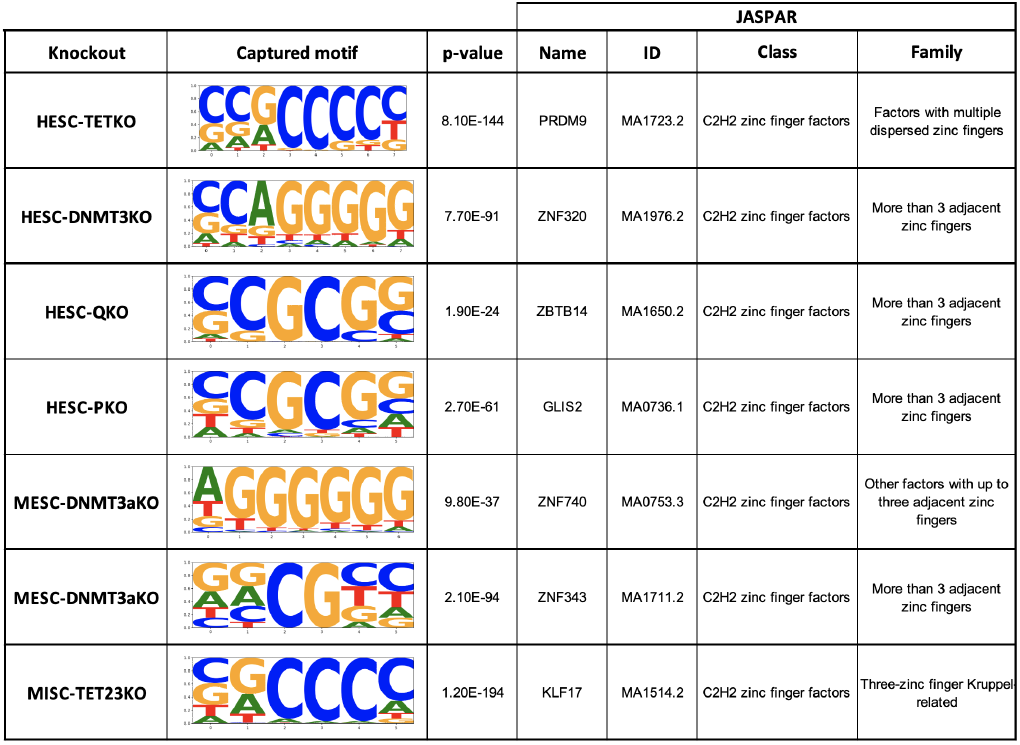
Sequence motifs (extracted from the attention layer of L-MAP) that achieved the lowest p-values in each knockout dataset and the corresponding the best hits from the JASPAR motif database

The first JASPAR hit in Figure 3 is the binding site for the PRDM9 zinc finger, which controls the location and intensity of crossovers during meiosis in humans and mouse (43; 45; 46). Studies have shown that there is a link between PRDM9 activity and TET1 during meiosis in mice (44). The second hit is the motif associated with ZNF320, which influences the regulation of cell cycle and immune infiltration, underscoring its significance in the molecular pathways of hepatocellular carcinoma progression (47). The third hit is the binding site for ZBTB14, which is a key protein in Xenopus embryonic development, influencing neural induction and differentiation by modulating BMP and WNT signaling pathways (48). ZBTB14 is also known as a regulator that binds to non-methylated CpG islands, playing a crucial role in controlling gene expression associated with the 2-cell-like state (49). The fourth hit is the motif associate with GLIS2, which has been identified as a transcriptional activator and is implicated as an epigenetically defined biomarker of a pluripotent phenotype in human ESCs (50). The fifth hit is the binding site for ZNF740, which plays a crucial role in cell differentiation by modulating the expression of MEF2C and its target genes, influencing the transition of pluripotent stem cells into trophoblasts through its interaction with a specific genomic variation (52). The sixth hit is the motif associate with ZNF343, which is involved in early stages of human embryonic development and influences embryo quality and developmental potential (51). The last hit on Figure 3 is the binding site of KLF17 which plays a significant role in the establishment of naive pluripotency in human ESCs (53).

L-MAP’s high accuracy in the prediction of DMCs for the TET knockout samples can be leveraged for a deeper analysis of the related motifs. In Supplemental File 1, we collected the 20 motifs with the lowest p-values and searched the JASPAR database for corresponding transcription factor binding motifs. These transcription factors can be further analyzed for potential interaction with TET. Notably, CTCF had the highest occurrence in the JASPAR hits. The interaction between CTCF and TET is well studied (67; 75; 74; 71; 72; 73). We expect that the other transcription factor in this list are also interacting with TET, but most of them are unexplored in the literature. This is an opportunity for research in functional determinants of TET proteins.

DNMT3a and DNMT3b *de novo* DNA methyltransferases are known to have strong sequence preferences, particularly in the sequences surrounding the CpG dinucleotides (76; 77; 78; 79; 80; 81). To investigate which positions in the input window are more important for the classification, we extracted the L-MAP’s attention scores. Figure 10 and Figure 11 in the supplemental material show the attention scores within the input window for L-MAP on different data sets. Observe that L-MAP’s strongest attention are the position flanking the center cytosine. Also observe that the attention is much stronger for the flanking positions for the DNMT3 data sets compared to TET data sets, consistent with the literature.

### Predictions using sequence and methylation levels

Within the scope of data imputation, one could assume to have the methylation levels of some cytosines and want to predict differentially methylated cytosines for the missing data. To test the extent of which imputation would be possible, we modified the input to L-MAP to allow the use of nearby methylation levels for the wild type sample, the knockout sample, or both (in addition to the primary DNA sequence surrounding the cytosine of interest). Details about the architecture of this variant of L-MAP can be found in the Methods section.

Figure 4 illustrates the performance of L-MAP using various input combinations. Observe that providing the methylation levels of neighboring cytosines does not significant improve L-MAP’s accuracy. In fact, in four out of seven cases, L-MAP performed slightly better when neighboring cytosine methylation levels were not provided.

**Fig. 4.**
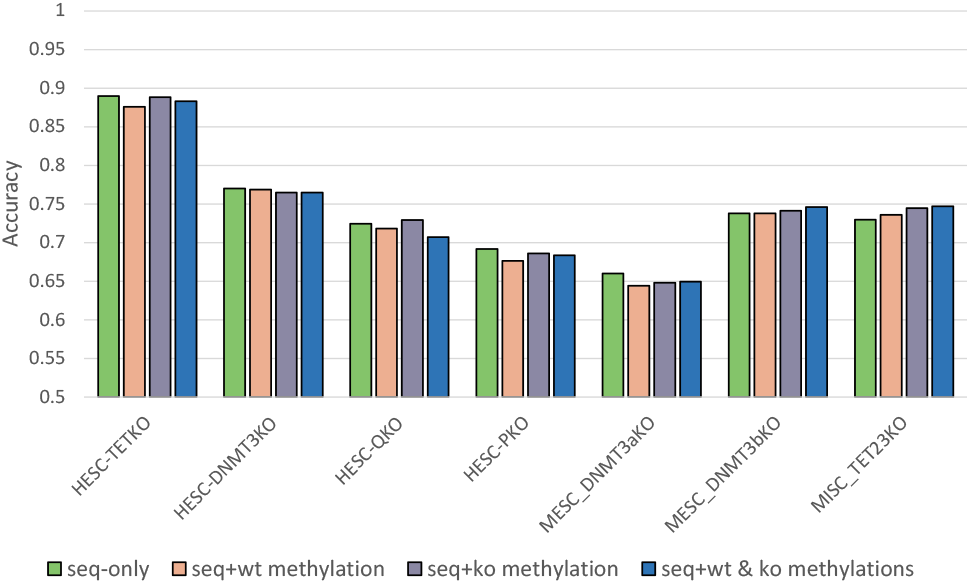
Effect of including neighboring cytosine methylation levels on L-MAP’s prediction accuracy for DMCs in seven knockout datasets

## Discussion

Here we introduced L-MAP, a large language model capable of predicting differentially methylated cytosines for TET and DNMT3 knockouts from the DNA sequence surrounding the cytosine of interest. Our findings highlight the potential of L-MAP to predict DMCs even when trained on different knockout datasets, with the exception of the model trained on the human TETKO dataset and tested on mouse ESC TET23KO and vice versa. This observation suggests distinct TET activity domains in ESCs between mouse and human species.

Furthermore, our study identified DNA sequence motifs associated with TET and DNMT3 activity in human ESC, mouse ESC, and mouse ISC, which were validated by comparing them to known motifs. Our work represents the first attempt in addressing this challenging problem, and it provides a tool to gain new insights into the role of TET and DNMT3 activity in cell processes, particularly during cell differentiation. The ability to predict DMCs and discover associated sequence motifs opens up opportunities for advancing our understanding of epigenetic regulation in various cellular processes.

## Key Points

- L-MAP is a large language model that can predict differentially methylated cytosines (DMCs) in human and mouse when trained on TET and DNMT3 knockout data sets
- L-MAP predicts DMCs with high accuracy exclusively based on the DNA sequence surrounding the cytosine of interest
- L-MAP can predict DMCs even when trained on different knockout data sets (human vs. mouse, or TET vs DNMT3)
- L-MAP can be used to discover new transcription factor associated with TET and DNMT3

## Methods

### Data sources and pre-processing

We used seven data sets from three different studies, namely (i) the human ESC knockout data sets generated by Charlton *et al*., who engineered several HUES8 embryonic stem cell lines using CRISPR-Cas9, producing variants with double, triple, quadruple, and quintuple genetic knockouts through the selective inactivation of DNMT3A, DNMT3B, TET1, TET2, and TET3 genes (2); (ii) the mouse knockout data sets generated by Gu *et al*. who analyzed the roles of DNMT3A and DNMT3B in DNA methylation within mouse ESCs following the loss of those enzymes (4); (iii) the mouse ISC knockout data sets produced by Ansari *et al*. who investigated the roles of TET2 and TET3 in the small intestine by generating double knockout mice (6). All the datasets are publicly available from NCBI. We note that in all these datasets, due to the choice of the protocol used to carry out bisufite-treated sequencing, only the methylation levels for the forward strand is available.

Given a pair of (wild type, knockout) data sets, we compared the difference in methylation levels for the same cytosine in the two experiments. We defined a cytosine to be differentially methylated (DMC) if the absolute value of the difference between the methylation level in the wild type and the methylation level in the knockout was at least 0.6, as proposed by Charlton *et al*. in (2). We only called DMC for cytosines that were covered by at least ten reads in both wild type and knockout experiments. Cytosines that were not covered by at least ten reads in either experiments were considered undetermined and ignored in our study.

### Training set design

We studied the effect of the size of the training set size on L-MAP’s accuracy in Supplemental Figure 7. Observe that L-MAP’s performance improves until the training set size reaches 100 thousand data points. Expanding the training set size further only increases the training time, without a significant benefit in the accuracy. Based on this analysis, for each experiment in our study, we sampled 50 thousand cytosines (uniformly at random) from all genome-wide cytosines that were differentially methylated, and another 50 thousand cytosines (uniformly at random) from all genome-wide cytosines that were not differentially methylated. We evaluated L-MAP’s performance for various choices of the input window sizes on DNMT3 and TET knockout datasets in Supplemental Figure 4. Based on this analysis we selected 512 bp centered around the cytosine of interest, as it yielded the best results among the tested sizes. We observe that 512 bp is the longest possible input that DNABERT allows. The sample containing 100 thousand sequences was divided into training set (90%) and test set (10%) uniformly at random. The label of each sequence was binary, indicating whether the center cytosine was differentially methylated or not.

### Classifiers

The architecture of L-MAP combines DNABERT (10) with a fully connected neural network as shown in Figure 1-A. In Supplemental Figure 6 we assessed the accuracy of other Transformer-based models. We selected DNABERT because it achieved the best performance on the TET knockout dataset. The input sequence was first tokenized in overlapping 6-mers. In Supplemental Figure 3 we tested various sizes for the tokens on the DNMT3 dataset, and *k* = 6 produced the best performance. DNABERT’s output layer was used as input to a fully connected neural network consisting of three layers with 128, 24, and 2 nodes respectively. Each layer used a dropout rate of 0.5 and employed the ReLU activation function, with the exception of the final layer, which utilized softmax as the activation function. The model was trained utilizing the Adam optimizer, with a learning rate of 1e-5, and employed the binary class entropy as the loss function.

In the experiments that used neighboring cytosine methylation levels, the embedding produced by DNABERT was concatenated with the vector(s) representing the methylation levels from either wild-type or knockout datasets (or both). This additional vector was -1 in all positions, except for the positions of neighboring cytosines with sufficient read coverage, where the known methylation level of the cytosine was used.

### Motif Analysis

We first obtained a random set of 10,000 genomic sequences of length 512 bp, where half of them were the context sequence surrounding a DMC, while the other half surrounded a non-DMC. We processed these sequences through DNABERT, then extracted the weights from DNABERT’s attention layer. We used the weights to identify high-attention regions, using the DNABERT motif-finding tool. For each of these regions, we extracted the corresponding DNA sequences from the original DNA sequence dataset, resulting in two distinct sets of DNA sequences for positive and negative samples. Then, we employed STREME (41) to identify motifs (and their p-values) that were enriched in the positive set and depleted in the negative set, using parameters minw=6, maxw=12, and nmotifs=100. The position weight matrices of the three motifs with the lowest p-values were matched against known motifs using JASPAR (38).

## Data access

All the datasets used in this study are publicly available from NCBI. The datasets accessions are GSE126958, GSE100956, and GSE200227. Bisulfite-treated Illumina reads were obtained from NCBI SRA, accessions SRR8611939, SRR6894127, and SRR18645747. L-MAP is available athttps://github.com/ucrbioinfo/dmc_prediction

### Abbreviations

5mC: = 5-methylcytosine
DMC: = differentially methylated cytosines LLM = large language model
L-MAP: = language model-based methyltransferases activity predictor
AUC: = area under the curve
TET: = ten eleven translocation (enzyme) DNMT = DNA methyltransferase (enzyme) ESC = embryonic stem cells
ISC: = intestinal stem cells

## Acknowledgements

The authors wish to thank Daniel Koenig (UC Riverside) and Jikui Song (UC Riverside) for earlier discussions on this project.

## Authors’ contributions

SS and SL conceptualized the project. SS designed L-MAP. SS carried out the experiments. SS and SL wrote the manuscript.

**Saleh Sereshki** is a Ph.D. candidate at the University of California, Riverside. He began his Ph.D. in Computer Science in 2019. His research focuses on applying machine learning techniques to genetic and epigenetic data.

**Stefano Lonardi** is Professor and Vice Chair of the Department of Computer Science and Engineering at the University of California, Riverside. His research interests include computational biology, bioinformatics, data mining, and epigenetics.

