## supplemental for "Predicting Differentially Methylated Cytosines in TET and DNMT3 Knockout Mutants via a Large Language Model": Sequence_level_TETs_and_DNMT3s_supp.pdf

### Supplementary material

Saleh Sereshki and Stefano Lonardi

### Supplementary Figures

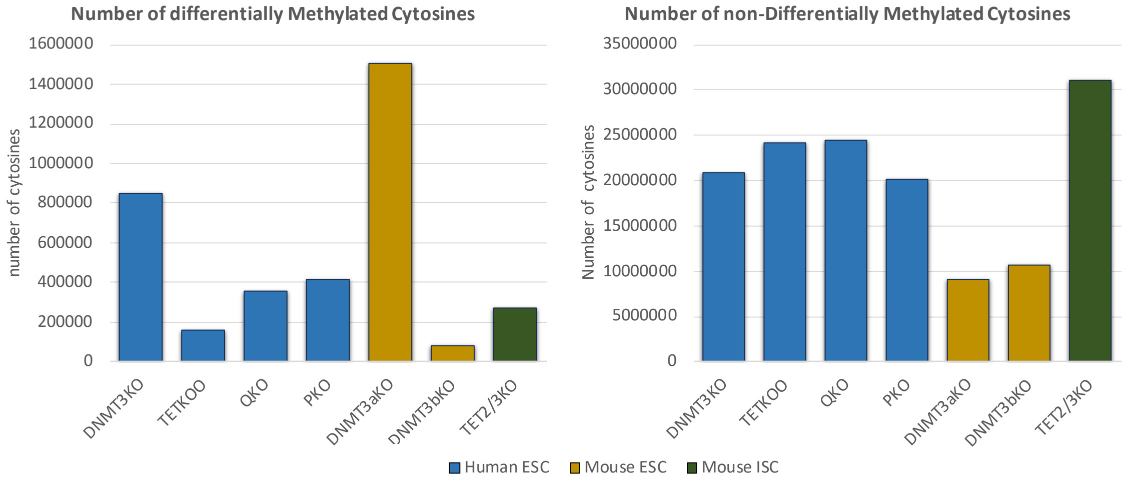


Supplementary Figure 1: (LEFT) the number of cytosines that are defined as "differentially methylated", i.e., cytosines that are covered by at least ten Illumina reads, and that have a methylation level in the knockout dataset that differs by at least 0.6 compared to the wild type; (RIGHT) the number of non-differential methylation in the seven datasets used in this study

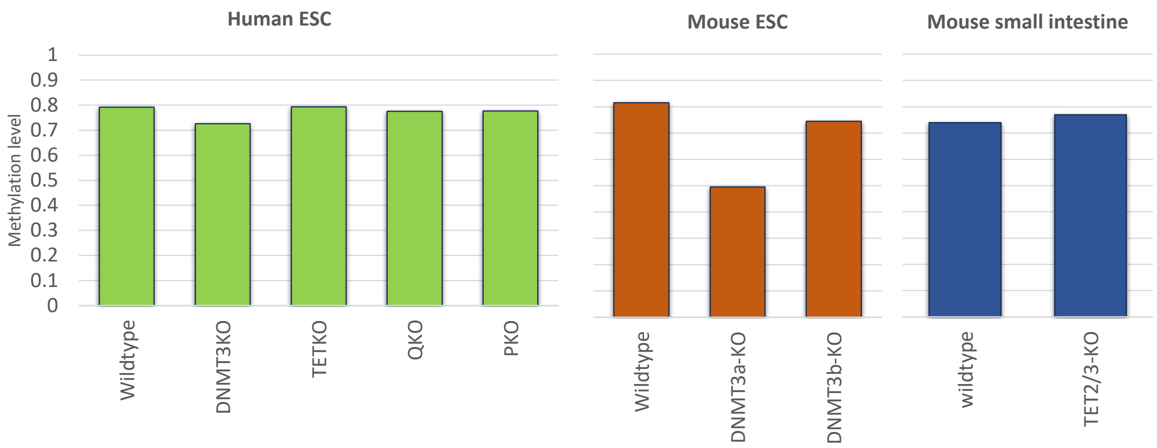


Supplementary Figure 2: The genome-wide average methylation level of cytosines across the seven knock-out and three wild type datasets

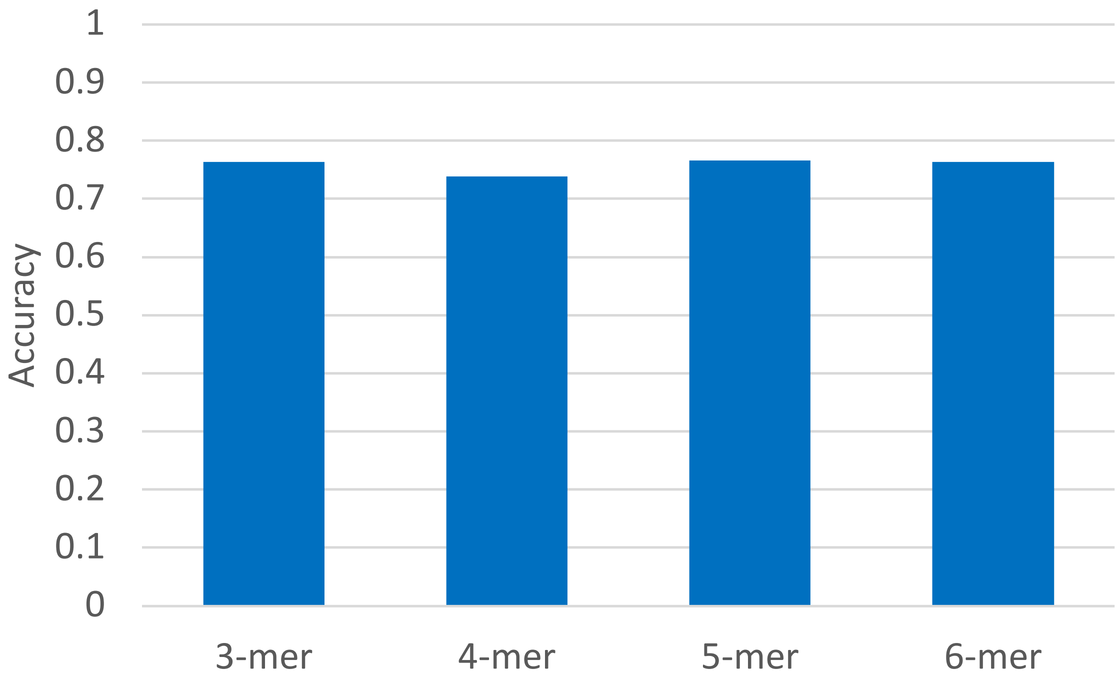


Supplementary Figure 3: L-MAP's performance in predicting differentially methylated cytosines in HESC cells with DNMT3 knockout, for various choices of the tokenization hyper-parameter $k$ in DNABERT

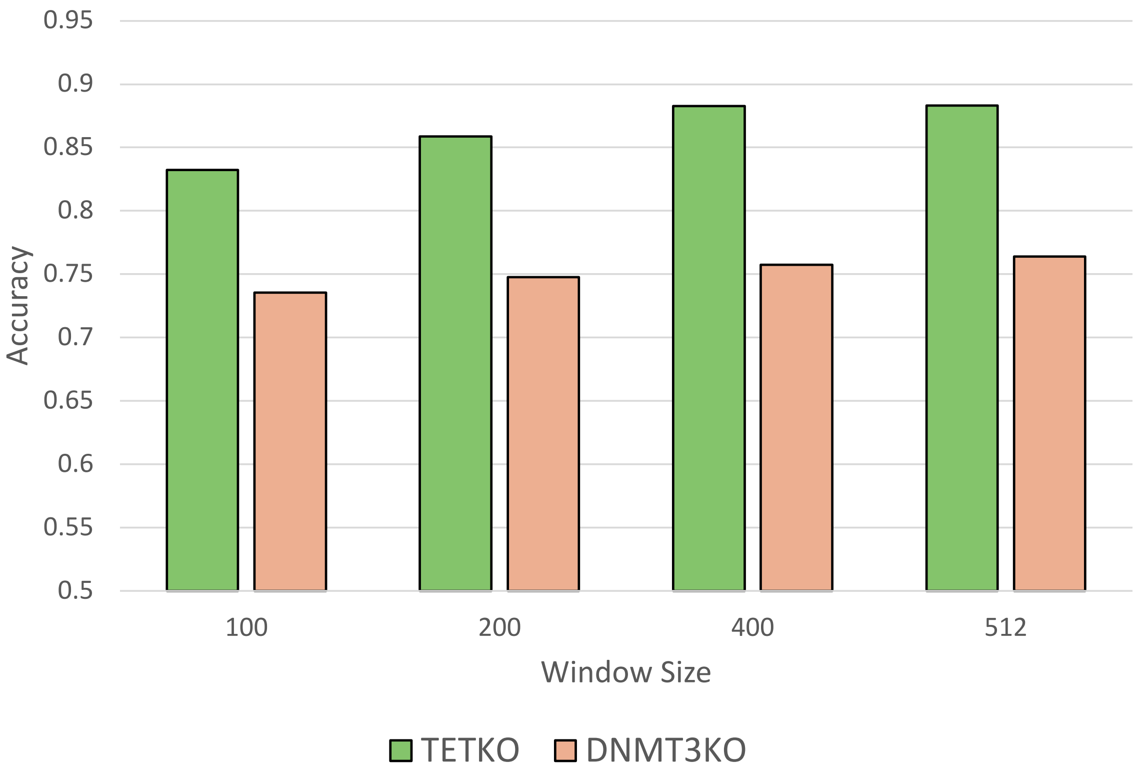


Supplementary Figure 4: Accuracy of L-MAP as a function of context sequence window size

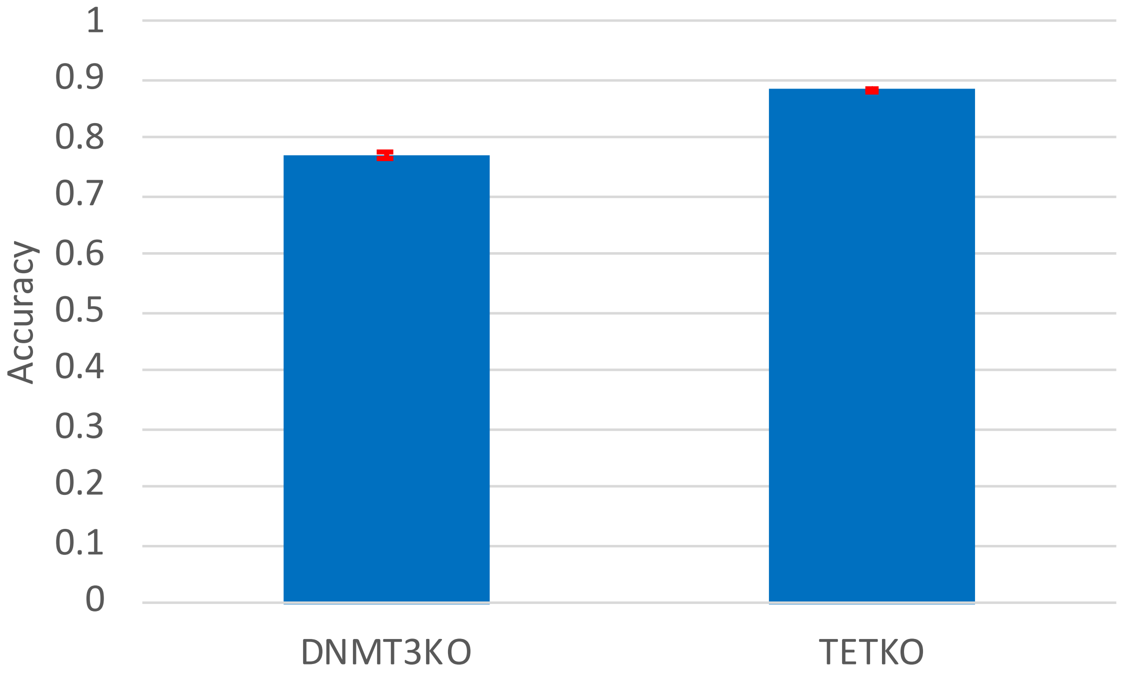


Supplementary Figure 5: Average and standard deviation of L-MAP's accuracy on the prediction of human DNMT3 and TET knockout over five experiments; in each experiment, L-MAP was trained on 90% of the cytosines (sampled uniformly at random from all the available cytosines), and tested on the remaining 10%

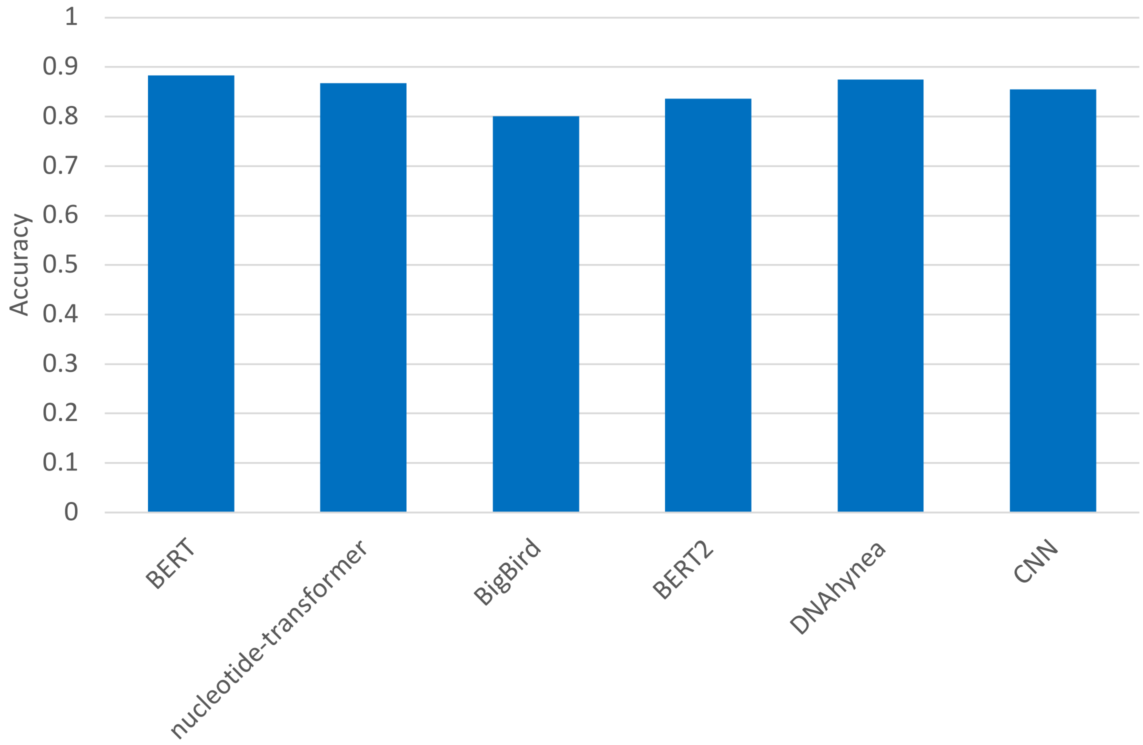


Supplementary Figure 6: Comparing the prediction accuracy of various large language models and a convolutional neural network (CNN) on the TET knockout dataset.

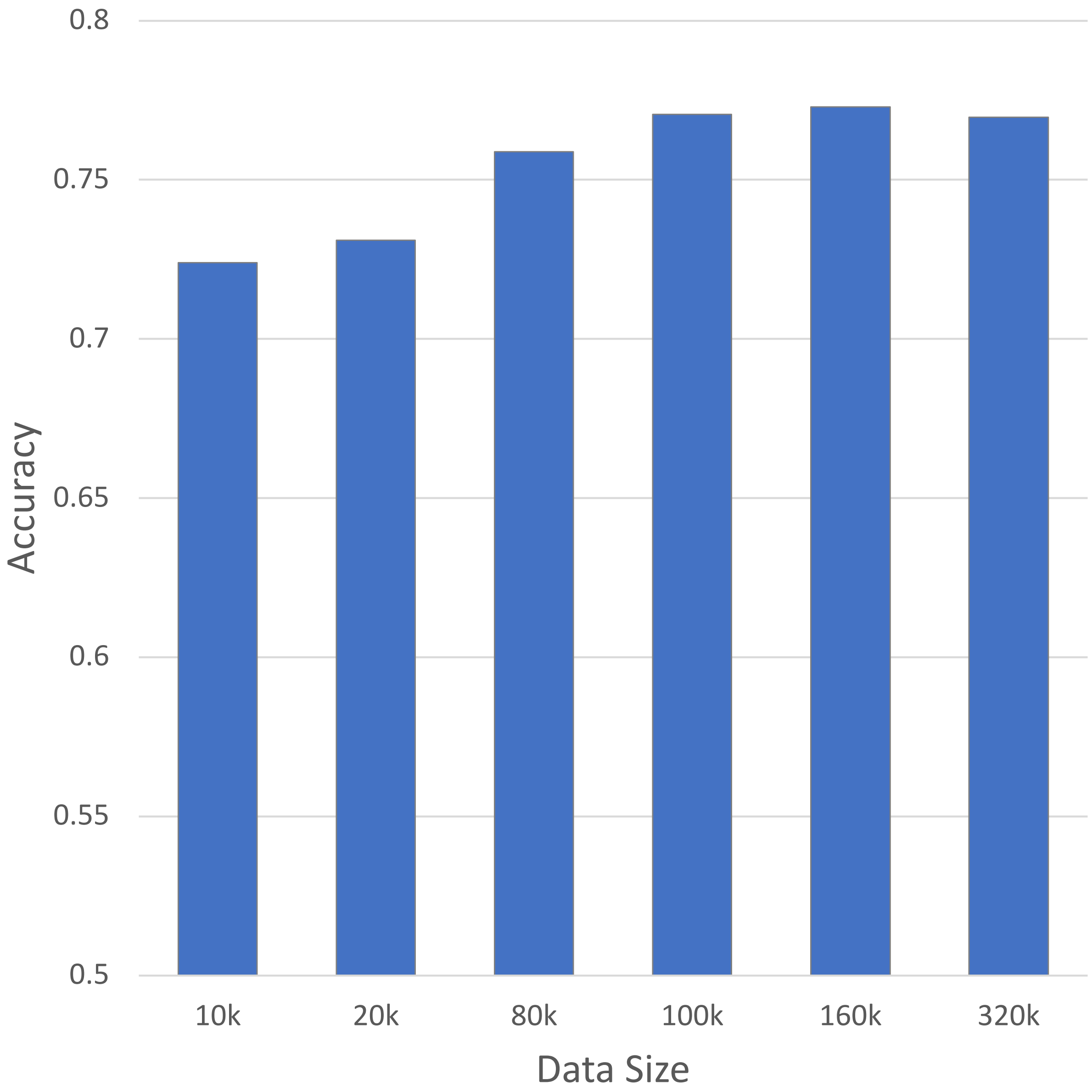


Supplementary Figure 7: L-MAP's prediction accuracy on the human DNMT3 knockout data for different choices of the training set size (half of these data points were differentially methylated, the other half was not)

| | | | JASPAR | | | |
|---|---|---|---|---|---|---|
| **Knockout** | **Captured motif** | **p-value** | **Name** | **ID** | **Class** | **Family** |
| **HESC-TETKO** | 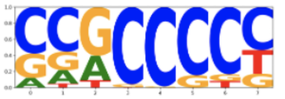 | 8.10E-144 | PRDM9 | MA1723.2 | C2H2 zinc finger factors | Factors with multiple dispersed zinc fingers |
| | 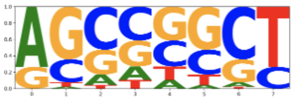 | 2.50E-38 | CTCF | MA1929.2 | C2H2 zinc finger factors | More than 3 adjacent zinc fingers |
| | 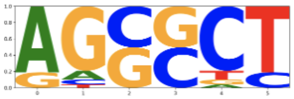 | 6.70E-13 | ZFP14 | MA1972.1 | C2H2 zinc finger factors | Factors with multiple dispersed zinc fingers |
| **HESC-DNMT3KO** | 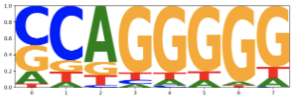 | 7.70E-91 | ZNF320 | MA1976.2 | C2H2 zinc finger factors | More than 3 adjacent zinc fingers |
| | 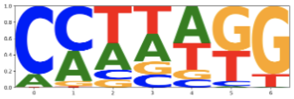 | 1.60E-11 | RARA::RXRG | MA1149.2 | Nuclear receptors with C4 zinc fingers::Nuclear receptors with C4 zinc fingers | Thyroid hormone receptor-related factors (NR1)::RXR-related receptors (NR2) |
| | 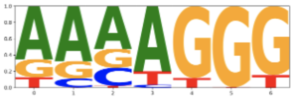 | 2.60E-06 | KLF13 | MA0657.2 | C2H2 zinc finger factors | Three-zinc finger Kruppel-related |
| **HESC-QKO** | 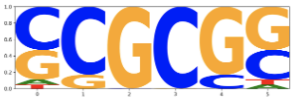 | 1.90E-24 | ZBTB14 | MA1650.2 | C2H2 zinc finger factors | More than 3 adjacent zinc fingers |
| | 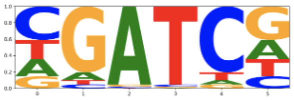 | 1.50E-18 | CUX2 | MA0755.2 | Homeo domain factors | HD-CUT |
| | 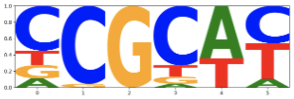 | 5.90E-08 | ZNF610 | MA1713.2 | C2H2 zinc finger factors | More than 3 adjacent zinc fingers |
| **HESC-PKO** | 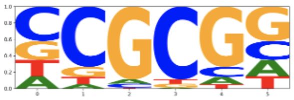 | 2.70E-61 | GLIS2 | MA0736.1 | C2H2 zinc finger factors | More than 3 adjacent zinc fingers |
| | 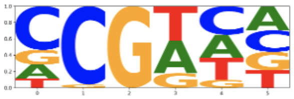 | 1.20E-32 | CTCF | MA1929.2 | C2H2 zinc finger factors | More than 3 adjacent zinc fingers |
| | 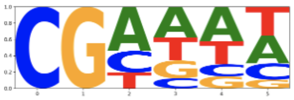 | 1.60E-06 | IRF5 | MA1420.1 | Tryptophan cluster factors | Interferon-regulatory factors |

Supplementary Figure 8: The three most statistically significant motifs extracted from L-MAP for each human knockout dataset and the corresponding the best hit from the JASPAR motif dataset

| | | | JASPAR | | | |
|---|---|---|---|---|---|---|
| **Knockout** | **Captured motif** | **p-value** | **Name** | **ID** | **Class** | **Family** |
| **MESC-DNMT3aKO** | 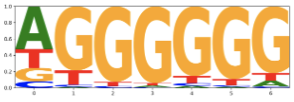 | 9.80E-37 | ZNF740 | MA0753.3 | C2H2 zinc finger factors | Other factors with up to three adjacent zinc fingers |
| | 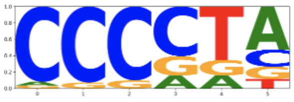 | 1.80E-05 | ZNF701 | MA1987.2 | C2H2 zinc finger factors | More than 3 adjacent zinc fingers |
| | 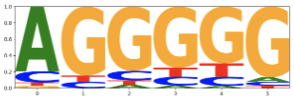 | 3.30E-05 | CTCF | MA1929.2 | C2H2 zinc finger factors | More than 3 adjacent zinc fingers |
| **MESC-DNMT3aKO** | 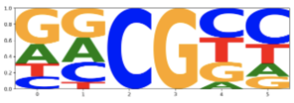 | 2.10E-94 | ZNF343 | MA1711.2 | C2H2 zinc finger factors | More than 3 adjacent zinc fingers |
| | 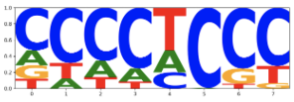 | 1.70E-46 | KLF5 | MA0599.1 | C2H2 zinc finger factors | Three-zinc finger Kruppel-related |
| | 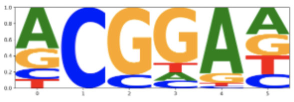 | 1.10E-21 | ETV5::FIGLA | MA1945.2 | Tryptophan cluster factors::Basic helix-loop-helix factors (bHLH) | Ets-related::Tal-related |
| **MISC-TET23KO** | 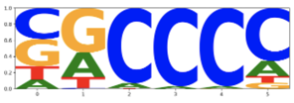 | 1.20E-194 | KLF17 | MA1514.2 | C2H2 zinc finger factors | Three-zinc finger Kruppel-related |
| | 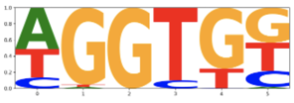 | 2.10E-31 | TBX19 | MA0804.2 | T-Box factors | Brachyury-related factors |
| | 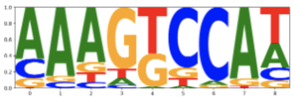 | 2.70E-28 | CTCF | MA1930.2 | C2H2 zinc finger factors | More than 3 adjacent zinc fingers |

Supplementary Figure 9: The three most statistically significant motifs extracted from L-MAP for each mouse knockout dataset and the corresponding the best hit from the JASPAR motif dataset

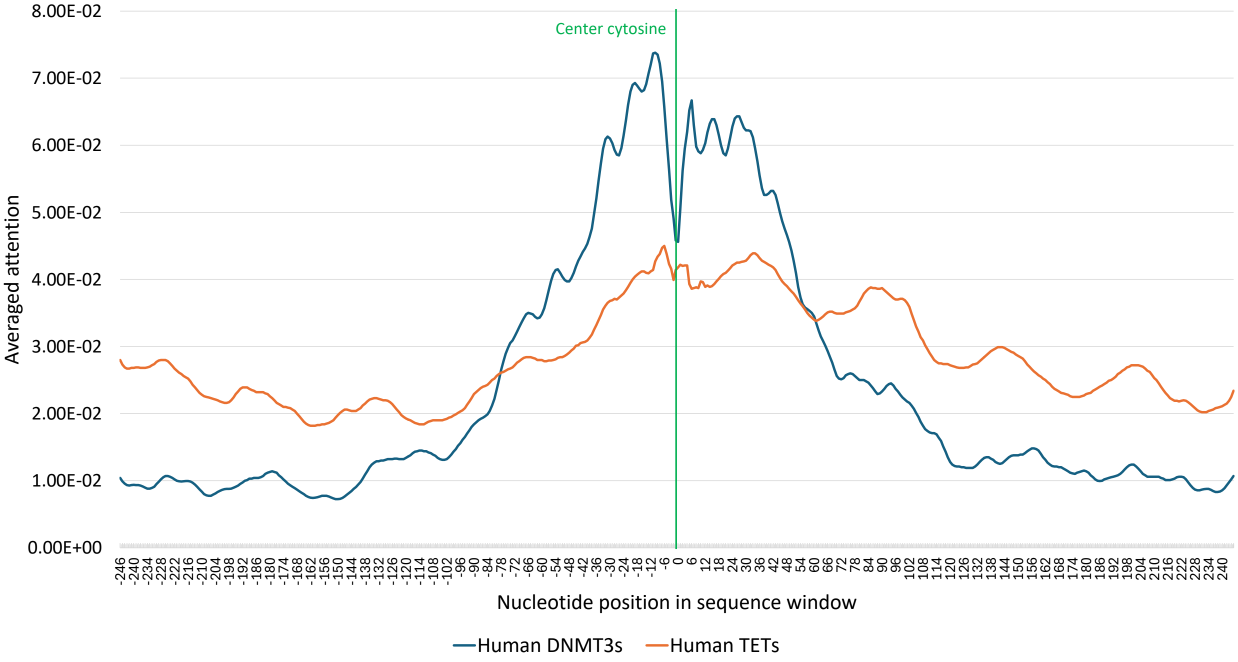


Supplementary Figure 10: Positional attention scores for L-MAP in the input window for human data sets

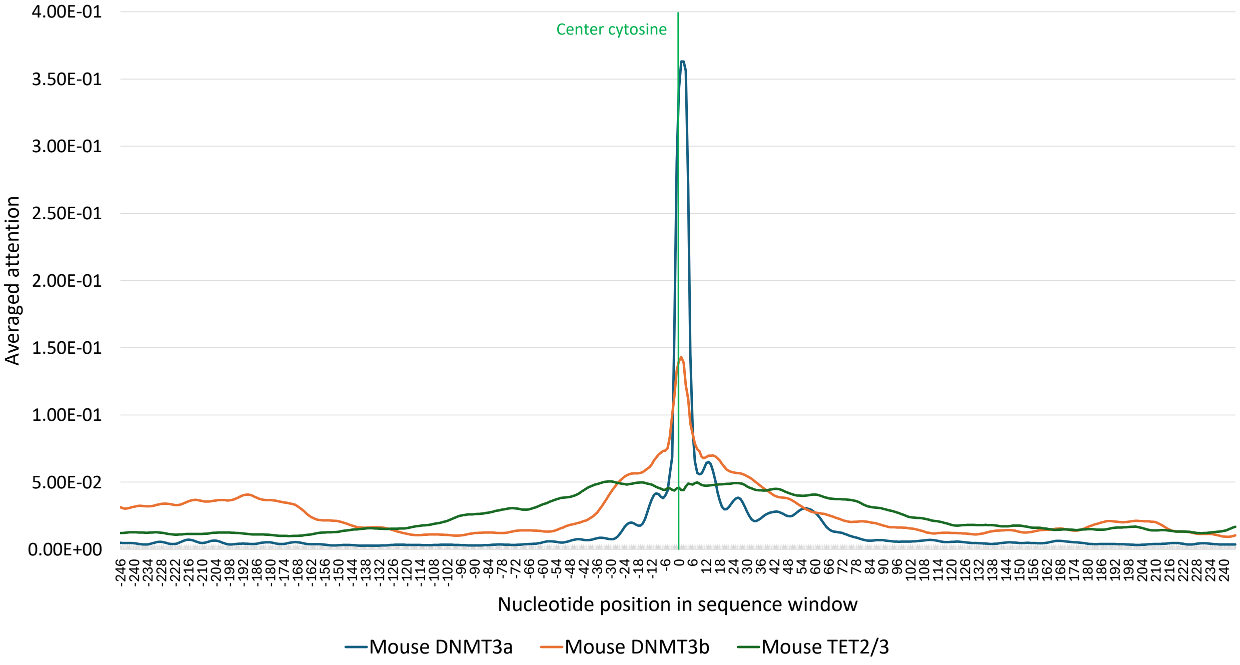


Supplementary Figure 11: Positional attention scores for L-MAP in the input window for mouse data sets
